# A novel structure-based approach for identification of vertebrate susceptibility to SARS-CoV-2: implications for future surveillance programmes

**DOI:** 10.1101/2022.01.10.475752

**Authors:** Rahul Kaushik, Naveen Kumar, Kam Y. J. Zhang, Pratiksha Srivastava, Sandeep Bhatia, Yashpal Singh Malik

## Abstract

Understanding the origin of severe acute respiratory syndrome coronavirus 2 (SARS-CoV-2) has been a highly debatable and unsolved challenge for the scientific communities across the world. A key to dissect the susceptibility profiles of animal species to SARS-CoV-2 is to understand how virus enters into the cells. The interaction of SARS-CoV-2 ligands (RBD on spike protein) with its host cell receptor, angiotensin-converting enzyme 2 (ACE2), is a critical determinant of host range and cross-species transmission. In this study, we developed and implemented a rigorous computational approach for predicting binding affinity between 299 ACE2 orthologs from diverse vertebrate species and the SARS-CoV-2 spike protein. The findings show that the spike protein of SARS-CoV-2 can bind to many vertebrate species carrying evolutionary divergent ACE2, implying a broad host range at the virus entry level, which may contribute to cross-species transmission and further viral evolution. Additionally, the present study facilitated the identification of genetic determinants that may differentiate susceptible from the resistant host species based on the conservation of ACE2-spike protein interacting residues in vertebrate host species known to facilitate SARS-CoV-2 infection; however, these genetic determinants warrant *in vivo* experimental confirmation. The molecular interactions associated with varied binding affinity of distinct ACE2 isoforms in a specific bat species were identified using protein structure analysis, implying the existence of diversified susceptibility of bat species to SARS-CoV-2. The findings from current study highlight the importance of intensive surveillance programs aimed at identifying susceptible hosts, particularly those with the potential to transmit zoonotic pathogens, in order to prevent future outbreaks.

## 1. Introduction

Detection of closely related viruses to that of Severe Acute Respiratory Syndrome-Coronavirus-2 (SARS-CoV-2) from animal sources is the most conclusive approach for identifying the zoonotic origins of SARS-CoV-2. Multiple evidences have emerged supporting immediate ancestor of SARS-CoV-2 is most likely a bat species (Kumar et al., 2021; Lytras et al., 2021; Malik et al., 2020; Zhou et al., 2021; Zhou et al., 2020), including the role of the intermediate host in transmission of SARS-CoV-2 to humans (Zhang et al., 2020). Although RaTG13, sampled from a *Rhinolophus affinis* bat in Yunnan, China (Zhou et al., 2021), has the highest nucleotide similarity to SARS-CoV-2, RaTG13 cannot be the recent progenitor of SARS-CoV-2 because it lacks evolutionary signatures possessed by SARS-CoV-2 (Kumar et al., 2021). After the emergence of SARS-CoV-2, three more bat viruses identified - RpYN06, RmYN02, and PrC31-that have a high nucleotide similarity in most of the virus genome, particularly ORF1b, implying that these viruses share a more recent common ancestor with SARS-CoV-2 (Li et al., 2021; Lytras et al., 2021; Zhou et al., 2021). Collectively, these studies demonstrate the zoonotic origin of SARS-CoV-2. However, to date, no bat reservoirs or intermediate animal hosts for SARS-CoV-2 have been identified. Therefore, it still remains unresolved about the spillover events leading to transmission of SARS-CoV-2 directly from bats to the humans or through an intermediate host.

The limited information about potential reservoir hosts, as well as the risk to wildlife and livestock, necessitate an immediate need of thorough investigation. Multiple SARS-CoV-2 experimental investigations and natural infection’s observations have demonstrated the variable susceptibility of vertebrate host species (such as mink, tigers, cats, gorillas, dogs, raccoon dogs, white tailed deer, rabbit, and ferrets) to SARS-CoV-2 infection (WHO, 2021; Cool et al., 2021; Malik et al., 2021a, 2021b; Mishra et al., 2021). Recent outbreaks in minks have linked SARS-CoV-2 transmission back to humans (reverse zoonosis) and to other animals (van Aart et al., 2021). However, except mink, there is no concrete evidence that other vertebrate host species are either spreaders of SARS-CoV-2 to humans (reverse zoonosis) or reservoir hosts. In addition, susceptibility of the majority of animal species that come into close contact with humans is unknown.

A key to dissect the susceptibility profiles of animal species to SARS-CoV-2 is to understand how virus enters into the host cells. SARS-CoV-2 spike protein initially binds to its receptor, angiotensin-converting enzyme 2 (ACE2), via its receptor-binding domain (RBD), before being proteolytically activated by proteases and performing its activity (Shang et al., 2020). Theoretically, the presence of ACE2 receptor in any host species makes them susceptible to SARS-CoV-2 infection, but this is not the case, even in animal species with significant ACE2 sequence similarity to human ACE2 (hACE2) (Liu et al., 2021). Therefore, a comprehensive understanding of the ACE2 diversity across the vertebrate species, combined with the protein-protein interactions at ACE2-RBD interface could facilitate some novel insights into the SARS-CoV-2 susceptibility in different vertebrate species.

In this study, we not only developed and implemented a rigorous computational pipeline for predicting 299 vertebrate ACE2 binding affinities (via dissociation constant) with the spike protein of SARS-CoV-2, but also demonstrated that dissociation constant is a better predictor of species susceptibility by benchmarking it with experimental data. By finding the best metric for assessing ACE2-RBD interactions, the molecular interactions leading to a variable binding affinity of ACE2 isoforms in a particular bat species to the spike protein of SARS-CoV-2 are revealed. Furthermore, based on the comparison of key interacting ACE2 residues at the ACE2-RBD interface, the genetic determinants that could aid in differentiating the SARS-CoV-2 susceptible from the resistant species are identified. Overall, the current study identifies a broad host range of vertebrate species susceptible to SARS-CoV-2 for further experimental investigations, along with proposing a novel approach for assessing animal species’ susceptibility profiles for viruses of interest.

## 2. Methodology

### 2.1 ACE2 Protein Sequences

The protein sequences of ACE2 orthologs (n = 356) originating from vertebrates were downloaded from NCBI protein database (5 September, 2021) (O’Leary et al., 2016). Partial or identical protein sequences were removed; however, ACE2 isoforms of a particular species were included in the final dataset. Therefore, the final dataset comprised of unique 299 ACE2 orthologs from 253 species. The species-wise number of the ACE2 orthologs retrieved from the NCBI protein database are provided in Supplementary Table S1.

### 2.2 Evolutionary divergence of ACE2 orthologs across the vertebrates

The full set of ACE2 protein sequences originating from vertebrates were aligned using MAFFT v.7.475 (Katoh et al., 2019) and a phylogenetic tree was constructed using the Maximum Likelihood statistical method with the GTR-Gamma model in RAxML v. 8.2.12 (Stamatakis, 2014) to understand the evolutionary relationship among the distinct vertebrate classes. Furthermore, using the Maximum Composite Likelihood model, evolutionary diversity and divergence between and within the vertebrate classes were estimated (Tamura et al., 2004). The number of base substitutions per site between sequences were used to calculate their evolutionary distance. Furthermore, the evolutionary divergence within and between the groups is represented by the number of base substitutions per site from averaging over all sequence pairs within each group and between the groups, respectively. The mean evolutionary diversity for the entire population 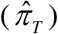 was calculated using equation (1).

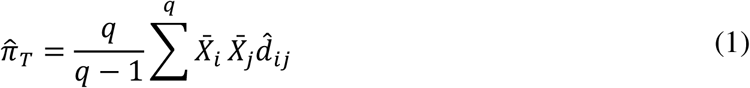

where, ‘*q*’ denotes the total number of alleles examined, 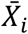 and 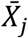 are the estimates of average frequency of the i^th^ and j^th^ alleles, respectively in the entire population, and 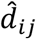 is the frequency of nucleotide substitutions per site between the i^th^ and j^th^ alleles. While the mean evolutionary diversity within subpopulations 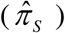 was calculated as shown in equation (2).

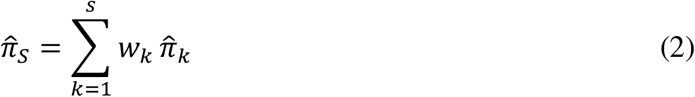

where, ‘*s*’ represents the subpopulations, the relative size of the k^th^ subpopulation is *w*_*k*_, and 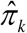 is the estimate of average nucleotide diversity in the k^th^ subpopulation. The inter-populational evolutionary diversity 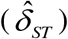 was calculated as per the equation (3)

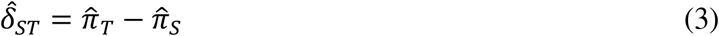

### 2.3 Model Structures for ACE2 Proteins

The recent accomplishments in the field of bioinformatics and computational biology, escorted to the development of some highly reliable de-novo, knowledge-based and hybrid methods for protein structure prediction that utilize the information inherent to the experimentally solved protein structures (Jumper et al., 2021; Yang et al., 2020; Yang et al., 2015; Bitencourt-Ferreira and de Azevedo, 2019; Kaushik and Jayaram 2016, Kaushik et al., 2018). We devised and executed a rigorous computational approach to model the 3-D structures for all of the ACE2 orthologs. Figure 1 depicts an overall workflow of the adopted computational pipeline in this study. For predicting model structures for each ACE2 protein, four state-of-the-art methods were used: online version AlphaFold2 (available through Colab-Notebook) (Jumper et al., 2021), standalone versions of Rosetta (*de novo* modeling) (Yang et al., 2020), I-Tasser (version 5.1) (*ab initio* and homology-based hybrid methodology) (Yang et al., 2015) and Modeller (version 10.1) (template-based modeling) (Bitencourt-Ferreira and de Azevedo, 2019), yielding twenty model structures (4 sets of 5 models from each method). The top scoring model structure was chosen from each set of model structures by using a consensus approach for protein structure quality assessment by implementing ProTSAV (Singh et al., 2016), ProFitFun (Kaushik and Zhang, 2020; Kaushik and Zhang, 2021) and ModFold8 (McGuffin et al., 2021). The top scored model structure for each ACE2 protein was refined through a comprehensive protein structure refinement approach implemented in GalaxyRefine (Heo et al., 2013). The refined model structures were re-evaluated with afore mentioned quality assessment methods to select the best scoring model structure for each ACE2 protein. The comprehensive approaches for protein structure quality assessment and protein structure refinement confirmed their reliability and applicability for studying various protein-protein interactions of ACE2 from vertebrates with the receptor binding domain of spike protein of SARS-CoV-2.

**Figure 1.**
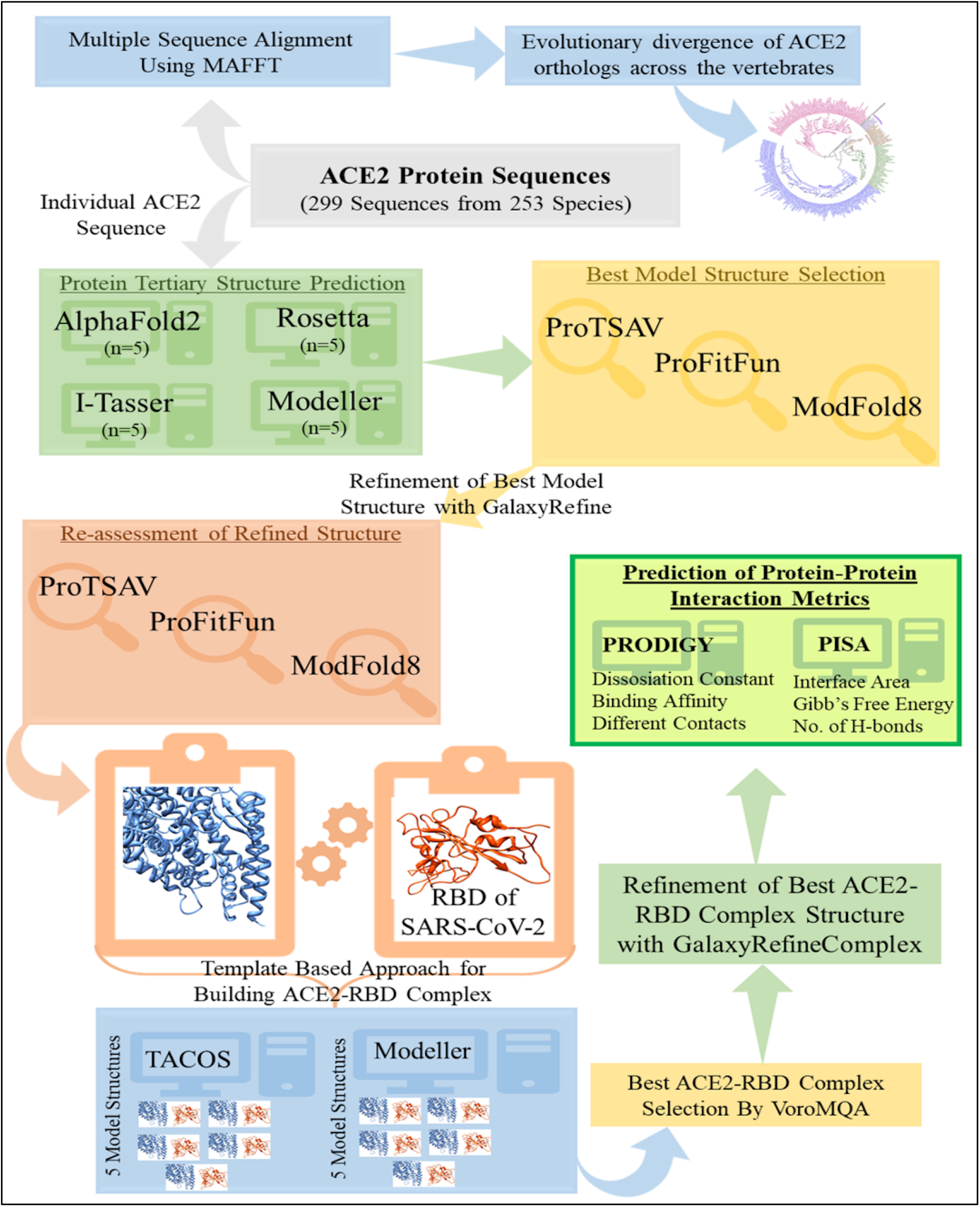
An overview of the computational methodologies used for predicting binding affinities of 299 vertebrate ACE2 proteins with RBD of SARS-CoV-2 spike protein in order to investigate the virus’s host range.

### 2.4 Template Based Approach for Building ACE2-RBD Complex

The reliability of template-based protein-protein complex predicting algorithms has improved as the number of experimentally determined protein-protein complexes has grown. The major challenge of modeling a protein-protein complex is finding an appropriate template that is well aligned with the target protein sequence. In general, the capacity to identify the interacting protein partners extracted by comparative modelling determines the efficacy of protein-protein docking in structural biology. Here, a template-based protein-protein docking was performed by implementing a standalone package of TACOS (Template-based Assembly of Complex Structures) (Szilagyi and Zhang, 2014) and the top ranked models of ACE2-RBD protein complex for each ACE2 protein were selected. Additionally, using ACE2-RBD complex of *Rhinolophus macrotis* (PDB ID: 7C8J), *Homo sapiens* (6M0J), *Canis lupus familaris* (7E3J), and *Falis catus* (7C8D) as reference structures, four more ACE2-RBD complexes for each ACE2 protein were predicted in the Modeller (version 10.1) (template-based modeling) (Bitencourt-Ferreira and de Azevedo, 2019). Finally, this led to 5 ACE2-RBD complexes for each ACE2 protein.

### 2.5 Refinement, Optimization of ACE2-RBD Complexes

The standalone version of GalaxyRefineComplex (Heo et al., 2016) was used to refine all of the ACE2-RBD docked complexes (n = 5 × 299). For each ACE2-RBD complex, five refined protein complexes were created, resulting in 25 ACE2-RBD complexes for each ACE2 protein. Using protein-protein complex quality assessment implemented by VoroMQA (Olechnovič and Venclovas, 2019), these protein complexes were ranked in order to choose the best refined and optimized ACE2-RBD complex for each ACE2 protein (n=299). The top-ranked ACE2-RBD protein complex for each ACE2 protein was utilized for carrying out further structural interaction analysis.

### 2.6 Benchmarking Binding Prediction Metrics on Dimeric Complexes

Different approaches for assessing protein-protein interactions in a protein complex yield different metrics for quantifying interaction strength. Some of these metrics include predicted binding affinity, predicted dissociation constant (Kd), change in Gibbs free energy (ΔG), binding energy, number of interactions (hydrophobic-hydrophobic, polar-polar, polar-hydrophobic, etc.), number of H-bonds, and number of di-sulfide bonds. Here, the PDBbind database was used to create a dataset of heterodimeric protein complexes (n = 400) in order to discover the best metric for assessing protein-protein interactions (Liu et al., 2015).

In this dataset of heterodimeric protein complexes, there are 154 complexes in total, each containing at least one enzymatic protein. Supplementary Table S2 has more information on these protein complexes. The experimental values of dissociation constants for these complexes, adopted from PDBbind Database (Liu et al., 2015), were used to compare the predicted protein-protein binding metrics. The standalone versions of PRODIGY (Xue et al., 2016) and PISA (Battle, 2016) were used to calculate various types of contacts (intermolecular, charged-charged, charged-polar, polar-polar, apolar-polar, and apolar-apolar), predicted binding affinity (Kcal/mol), predicted dissociation constant (Kd), interface area (Å), number of hydrogen bonds, salt bridges, disulfide bonds, and change in Gibbs free energy (ΔG). A correlation analysis of these parameters with experimental values of dissociation constants was performed to identify the most accurately predicted metric. It was observed that the predicted dissociation constant showed the best correlation with the experimental values. Considering this, the predicted dissociation constant was used as the main metric for quantifying the protein-protein interaction between ACE2-RBD complexes.

### 2.7 Interface and Binding Prediction of ACE2-RBD Complex

Using the standalone version of PRODIGY (Xue et al., 2016), a state-of-the-art method for predicting the binding in protein-protein complexes, the interacting interfaces of RBD and ACE2 were extracted from the top ACE2-RBD protein complex for each ACE2 protein and analyzed to calculate the dissociation constant. PRODIGY was also used to determine anticipated binding affinity for various types of contacts (intermolecular, charged-charged, charged-polar, polar-polar, apolar-polar, and apolar-apolar). Furthermore, the interface area, number of hydrogen bonds, salt bridges, and disulfide bonds, and change in Gibbs free energy (ΔG) were also estimated using the standalone version of PISA (Battle, 2016), implemented in CPP4 software suite (Supplementary Table S3). The predicted dissociation constants were utilized to quantify and build a better understanding of SARS-CoV-2’s host preferences.

### 2.8 Statistical analysis

For statistical analysis, GraphPad Prism 7.01 was used (GraphPad Software, San Diego, CA). We used multiple t-tests with Holm-Sidak method adjustments to assess the variations in evolutionary distances across the vertebrate classes. A p-value of less than 0.01 was considered statistically significant. GraphPad Prism 7.01 software was used to create all of the graphs.

## 3. Results

### 3.1 ACE2 protein’s evolutionary diversity across a wide range of vertebrate species

The protein sequences of 299 ACE2 orthologs downloaded from the NCBI protein database were analyzed and classified into six vertebrate taxonomic classes: 86 *Actinopterygii* (bony fishes), 6 *Amphibia* (amphibians), 43 *Aves* (birds), 2 *Chondrichthyes* (cartilaginous fishes), 143 *Mammalia* (mammals), and 19 *Reptilia* (reptiles), and subsequently estimated their evolutionary diversity and divergence (Figure 2). All the ACE2 protein sequences have a mean evolutionary diversity (d±SE) of 0.43±0.02, showing a huge diversity among these protein sequences. Besides, the mean evolutionary diversity of ACE2 protein sequences between and within the classes was 0.20±0.01 and 0.22±0.01, respectively. In addition, the average evolutionary divergence (d±SE) of ACE2 protein sequences among the six vertebrate taxonomic classes ranged from 0.27±0.01 to 0.13±0.009, where *Aves* had the least average evolutionary divergence among all the vertebrate classes (P < 0.0001) (Figure 3A). Furthermore, ACE2 protein sequences from *Mammalia* have a statistically higher mean evolutionary divergence than ACE2 from *Aves* (P < 0.0001), and lower as compared to ACE2 from *Actinopterygii* (P = 0.0006) (Figure 3B). Nevertheless, the mean evolutionary divergence of ACE2 protein sequences from *Mammalia* is non-significantly related to *Amphibia, Reptilia*, and *Chondrichthyes* (P = 0.018-0.569).

**Figure 2.**
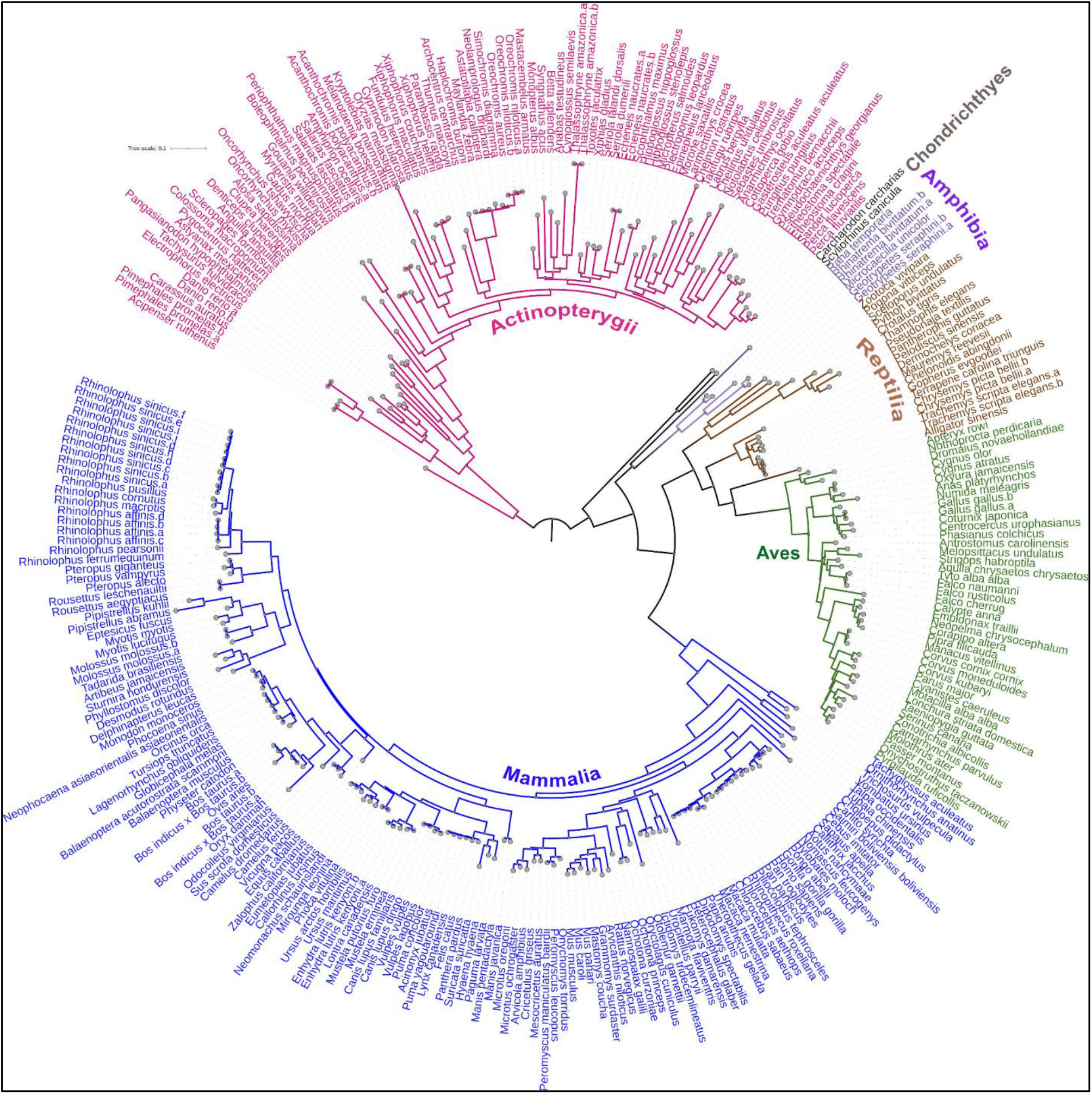
Maximum-likelihood (ML) tree showing the phylogenetic relationships of complete 299 ACE2 protein sequences derived from vertebrate species. The ACE2 protein sequences are classified into six vertebrate taxonomic classes; *Actinopterygii* (bony fishes), *Amphibia* (amphibians), *Aves* (birds), *Chondrichthyes* (cartilaginous fishes), *Mammalia* (mammals), and *Reptilia* (reptiles), and are colored differently.

**Figure 3.**
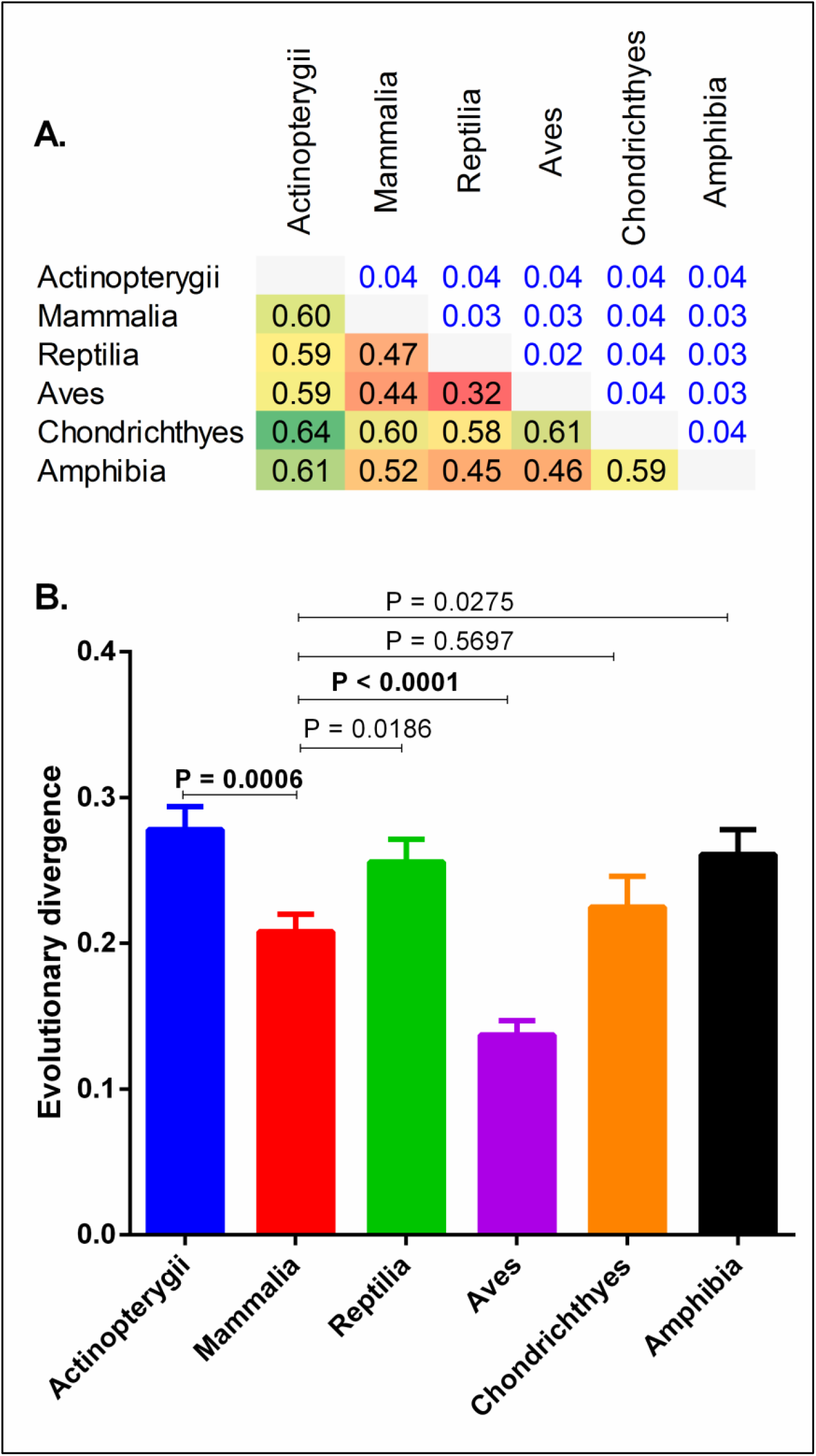
(**A**) shows a matrix showing the evolutionary divergence among the six vertebrate taxonomic classes. (**B**) shows a comparison of mean evolutionary divergence between the six vertebrate taxonomic classes. A *p*-value less than 0.01 was considered statistically significant.

In addition, the ACE2 protein sequences of *Homo sapiens* showed the highest sequence identity of 99.38% with that of *Hominidae* family members (*Gorilla gorilla gorilla, Pan paniscus*, and *Pan troglodytes)* followed by the *Hylobatidae* family members, with sequence identities of 98.76% and 98.29% (*Hylobates moloch* and *Nomascus leucogenys*, resectively), and *Cercopithecidae* family members (*Chlorocebus* sp., *Macaca* sp., *Papio Anubis, Piliocolobus tephrosceles, Rhinopithecus roxellana*, and *Theropithecus gelada*) with sequence identities of 96.05-96.86% (Supplementary Table S4). However, the ACE2 protein sequences of *Homo sapiens* showed poor sequence identities with the distinct ACE2 isoforms of *Rhinolophus* sp. (77.0-78.14%) and also with different species of bats, such as, *Pteropus* sp. (76.31-78.59%), *Pipistrellus* sp. (70.47-71.48%), *Myotis* sp. (77.46%), *Desmodus rotundus* (76.54%), and *Eptesicus fuscus* (77.92%). These findings show that the ACE2 protein sequences in *Mammalia* have a great deal of evolutionary diversity.

### 3.2 Prediction of vertebrate ACE2 orthologs binding affinities to spike protein of SARS-CoV-2

In order to discover the best metric for assessing ACE2-RBD interactions, the predicted dissociation constant (Kd), and predicted the change in Gibbs free energy (ΔG) were benchmarked against the experimental values (n = 400) retrieved from the PDBbind database (Liu et al., 2015). The benchmarking of these two matrices on the predicted and experimentally values derived from 400 protein complexes revealed that the results of predicted dissociation constant are highly correlated with experimental values (r = 0.858, P < 0.0001), while that of predicted change in Gibbs free energy (ΔG) is poorly correlated with the experimental values (r = 0.125, P < 0.012). Therefore, we first devised and conducted a rigorous computational approach to construct ACE2-RBD models for all of unique 299 ACE2 orthologs, and subsequently predicted dissociation constant (Kd) for assessing their interactions. Intriguingly, the results in the present study, based on the predicted dissociation constant, showed that the binding affinity of dACE2 with RBD is 4.15 times lower than that of hACE2, which is comparable (6.65 times lower) to an experimental study that calculated the binding affinity of dACE2/hACE2 to RBD using surface plasmon resonance (Zhang et al., 2021). Subsequently, for assessing ACE2-RBD interactions on the basis of predicted dissociation constant, the vertebrate species were categorized into four groups based on their propensity to bind to SARS-CoV-2: very high, high, medium, and low. The results for all the vertebrate species are provided in Supplementary Table S5. The 7 species predicted as very high consisted of 3 mammals (Southern elephant seal, Cat, and North American river otter), 3 reptiles (Chinese Alligator, Leatherback sea turtle, and Plateau fence lizard), and 1 bird species (White-ruffed manakin). Apart from this, ACE2 protein of 120 (16 bony fishes, 18 birds, 75 mammals, and 11 reptiles), 155 (61 bony fishes, 23 birds, 62 mammals, 5 reptiles, 3 amphibians, and 1 cartilaginous fish) and 17 species (10 bony fishes, 3 mammals, 3 amphibians and 1 cartilaginous fish), respectively were classified to have a high, medium, and low propensity to bind to SARS-CoV-2 (Supplementary Table S5).

Next, a comparison of the species’ predicted propensity (on the basis of predicted Kd) to bind to SARS-CoV-2 with the experimental proven susceptibility of infection (both the natural and experimental infection) was performed in 46 species (high, medium, low, and extremely low), as depicted in Figure 4. Intriguingly, all these 46 species were distributed across the very high, high and medium propensity categories predicted in our study. For example, it was found that 13 species designated as highly susceptible to SARS-CoV-2 infection on the basis of experimental studies fall across very high (n = 1, cat), high (n = 7, White-tailed deer, Human, Western lowland gorilla, Ferret, Prairie deer mouse, Golden hamster, and Mountain lion), and medium (n = 5, Leopard, Rhesus macaque, Common marmoset, American mink, and Egyptian rousette) propensity categories of our study, while one species (Rabbit) designated as medium susceptible to SARS-CoV-2 infection occupied medium propensity category of our study.

**Figure 4.**
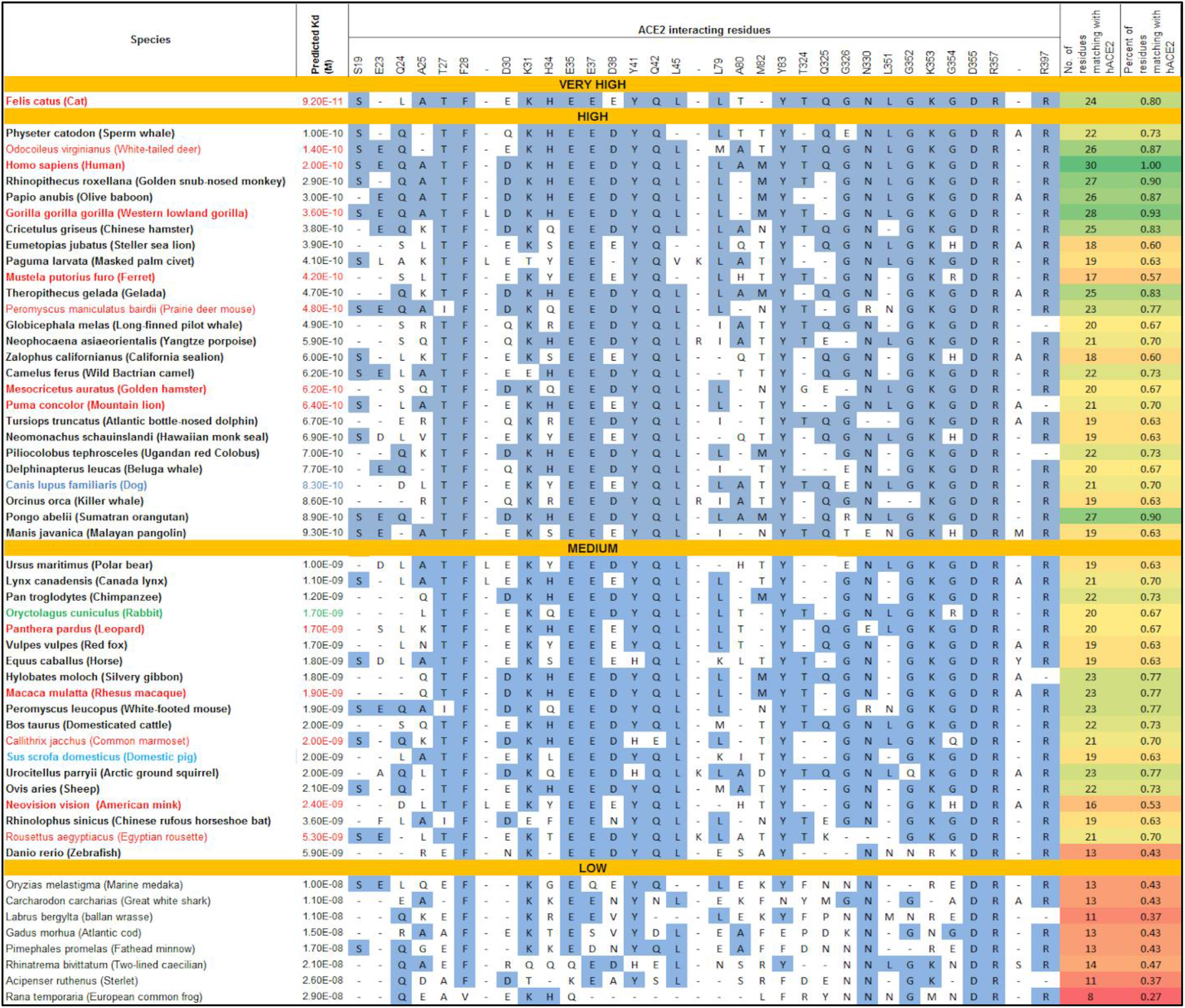
Comparison of predicted ACE2 binding affinity and their interacting residues among experimental proven susceptibility species. Species are sorted into very high, high, medium and low propensity using binding affinity scores. A comparison of ACE2 interacting residues at the ACE2-RBD interface with the fraction of interacting ACE2 residues similar to hACE2 is also shown. Experimentally proven high, medium, low and extremely low susceptibility species to SARS-CoV-2 are colored as red, green, blue, and sky blue, respectively. ACE2 interacting residues similar to hACE2 are highlighted.

### 3.3 Comparison of predicted ACE2 binding affinity and their interacting residues among experimental proven susceptibility species

We identified a total of 54 species of which 46 species were proven to support the SARS-CoV-2 infection either naturally or experimentally, while the rest of the 8 species predicted as resistant to SARS-CoV-2 infection were taken as control for comparative studies in the present study. Subsequently, we combined and compared their predicted ACE2 binding affinity (Kd) with the fraction of interacting ACE2 residues similar to hACE2, to gain insight into their differential propensity to bind to SARS-CoV-2 (Figure 4). The predicted dissociation constant is negatively correlated with the fraction of interacting ACE2 residues similar to hACE2 (r = −0.5997, P < 0.0001), implying that binding affinity of ACE2 to the spike protein of SARS-CoV-2 is associated with the number of interacting ACE2 residues similar to hACE2. Subsequently, we identified 30 amino acid residues in ACE2 interacting with the spike protein’s RBD of SARS-CoV-2 and these residues were examined for their similarity to the residues in hACE2. On the basis of number of residues identical to hACE2, the species susceptibility was classified into three categories (high, medium, and low); at least 24 residues identical to hACE2 for high (at least 24/30), 15-23 residues for medium (15 to 23/30), and less than 14 residues for low (less than 14/30) category. Nine species (Cat, White-tailed deer, Human, Golden snub-nosed monkey, Olive baboon, Western lowland gorilla, Chinese hamster, Gelada, and Sumatran orangutan) that were predicted to have a high propensity (based on the similarity of at least 24/30 residues with that of hACE2) also had very high to high propensity on the basis of predicted dissociation constant. However, comparing the ACE2 residues for their similarity to the residues in hACE2 alone does not truly reflect their propensity for SARS-CoV-2 because experimentally proven highly susceptible species to SARS-CoV-2 do not necessarily possess comparatively higher number of ACE2 interacting residues than hACE2, for example, Ferret (17/30), American mink (16/30), and Leopard (20/30). Of note, interpreting the susceptibility of species for SARS-CoV-2 should be done with caution and therefore, combining predicted ACE2 binding affinity (Kd) with the fraction of interacting ACE2 residues similar to hACE2 could yield more confidence in predicted propensity of species susceptibility to SARS-CoV-2 infection. The species predicted as low susceptibility to SARS-CoV-2 infection all carried ≤ 14/30 residues similar with that of hACE2 and also had a high Kd values (low binding affinity).

Previous studies had identified five critical ACE2 interacting residues based on their conservation in known susceptible species (31K/T, 35E/K, 38D/E, 82T/M/N, and 353K) (Shang et al., 2020; Wan et al., 2020). We carried out sequence alignment of ACE2 interacting residues of 54 species, and our analysis revealed that 31K/T, 35E/K, 38D/E, and 353K residues despite being highly conserved in high and medium susceptible species, are also observed in some of the predicted low susceptible or resistant species. In contrast, 82T/M/N residues are highly conserved and are restricted to high and medium susceptible species only. Based on the sequence alignment of ACE2 interacting residues, our analysis proposes six unique residues that could together help in differentiation of susceptible from the resistant species; Susceptible species (27T/I, 30D/E/Q, 82M/T/D/N, 326G/E/R/T, and 352G+353K+354G), resistant species (N326+N330), and 352G+353K+354H/R/Q for reduced susceptibility to SARS-CoV-2 infection.

Next, based on the predicted binding affinity (in terms of Kd), similarity score (ACE2 interacting residues similar to that of hACE2), and six unique residues proposed in this study as determinant of SARS-CoV-2 susceptibility together, 10 bat species (*Rhinolophus sinicus, R. affinis, R. macrotis, R. pearsonii, R. ferrumequinum, Myotis lucifugus, Artibeus jamaicensis, Rousettus aegyptiacus, Pteropus vampyrus*, and *Myotis myotis*) were predicted as medium susceptibility to SARS-CoV-2 infection, while the rest of 8 bat species were predicted to be resistant to SARS-CoV-2 (Supplementary Table S6). Intriguingly, of the identified 10 ACE2 isoforms in *Rhinolophus sinicus* and 4 ACE2 isoforms in *Rhinolophus affinis*, 2 each of them were predicted to have a high binding affinity with the spike protein of SARS-CoV-2.

### 3.4 Structural insights into differential ACE2 isoforms binding affinity

The interaction interface of hACE2, dACE2, rsACE2 (two isoforms), and raACE2 with RBD were analyzed separately from the refined ACE2-RBD complex using PRODIGY (Xue et al., 2016) and subsequently compared with hACE2-RBD interface to investigate the molecular basis of differential binding affinity of ACE2 across these species. The residues-wise atomic contacts of ACE2 with RBD for the selected vertebrate species are provided in Supplementary Table S7. We noted that 30 hACE2 residues formed a total of 79 atomic contacts with 25 RBD residues, of which 13 H-bonds and 2 salt bridges formed between 10 ACE2 (GLN 24, ASP 30, GLU 37, ASP 38, GLN 42, TYR 83, ASN 330, LYS 353, ASP 355, and ARG 357) and 8 RBD (LYS 417, GLY 446, TYR 449, ASN 487, TYR 489, THR 500, TYR 505, and GLY 502) residues. Of note, the total number of atomic contacts at the hACE2-RBD interface, including H-bonds and salt bridges are more than that at dACE2-RBD (78 atomic contacts, 12 H-bonds and 2 salt bridges), rs1ACE2-RBD (74 atomic contacts, 9 H-bonds and 1 salt bridge), rs10ACE2-RBD (70 atomic contacts, 8 H-bonds and 1 salt bridge), and raACE2-RBD (71 atomic contacts, 8 H-bonds and 1 salt bridge), which is consistent with our finding that hACE2 has higher predicted binding affinity as compared to these vertebrate species as depicted in Figure 5. Additionally, one isoform of rsACE2 (74 atomic contacts between 31 ACE2 and 24 RBD residues) made more atomic contacts including H-bonds with RBD residues as compared to another isoform of rsACE2 (70 atomic contacts between 26 ACE2 and 26 RBD residues), which support the presented finding that different isoforms of ACE2 in a particular bat species can have differential binding affinity with RBD. Intriguingly, engaging a small number of RBD residues (n = 25) by the virus in forming atomic contacts with a large number of ACE2 residues (n = 30) as noted at hACE2-RBD interface was also observed for one of the isoforms of rsACE2 and raACE2, but not in dACE2-RBD interface, possibly suggesting molecular interactions optimization at ACE2-RBD interface, and therefore, molecular signatures implying the bat origin of SARS-CoV-2 (Figure 5).

**Figure 5.**
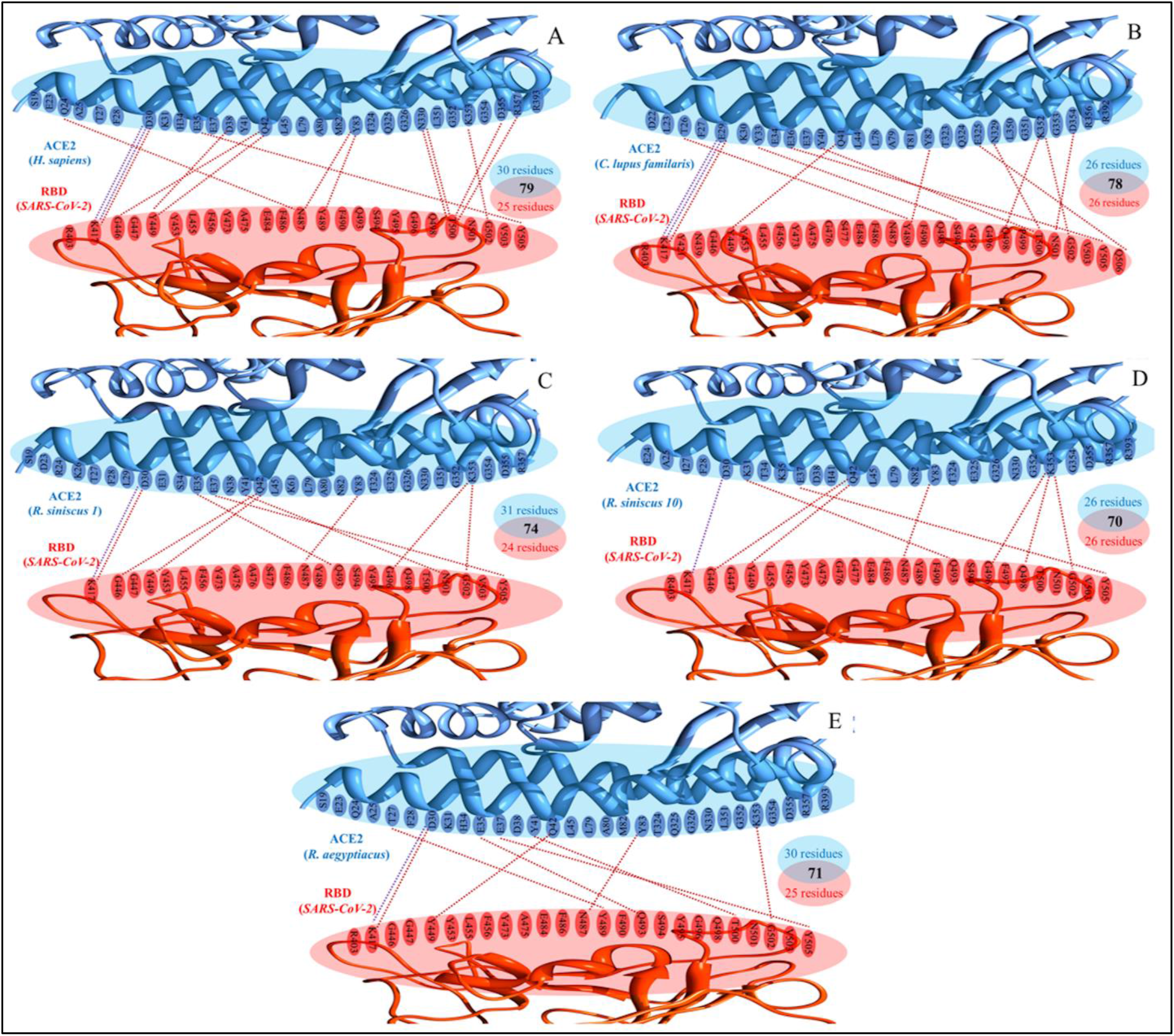
Structural insights into differential ACE2 binding affinities of selected vertebrate species to SARS-CoV-2 spike protein. The molecular interactions at the ACE2-RBD interface for **(A)** *Homo sapience*, **(B)** Canis lupus familaris, **(C)** *Rhinolophus sinicus* isoform 1 (GenBank accession no. 1883189465), **(D)** *Rhinolophus sinicus* isoform 10 (GenBank accession no. 1883189431), and **(E)** *Rousettus aegyptiacus* are shown. The red and blue dotted lines at the ACE2-RBD interface represent the hydrogen bonds salt bridges, respectively. The total number of interacting residues of ACE2 (blue) and RBD (red), including the total intermolecular contacts at ACE2-RBD interface are also shown.

## 4. Discussion

The interaction of SARS-CoV-2 with its host cell receptor is a critical determinant of host species susceptibility, tissue tropism, and viral pathogenesis. The RBD of SARS-CoV-2 spike protein recognizes the ACE2 receptor on the host cells, which allows the virus to enter the host cells (Shang et al., 2020a; Shang et al., 2020b). To explore the possible origin of SARS-CoV-2, species at risk, and species that could potentially serve as the intermediate hosts, we first presented a deeper understanding of ACE2 evolutionary diversity, followed by structural insights at the ACE2-RBD interface in a variety of vertebrate species.

Computational tools are the method of choice for examining the protein-protein interactions in a protein complex, especially when studying a big dataset, because many of the protein structures are yet to be solved experimentally. To forecast different species susceptibility to SARS-CoV-2, previous studies relied either on the comparison of twenty-five amino acids corresponding to the known SARS-CoV-2 Spike protein receptor binding residues for their similarity to the residues in human ACE2 or on the prediction of binding energies (Damas et al., 2020; Lan et al., 2020; Liu et al., 2021b; Shang et al., 2020a; Shang et al., 2020b; Sun et al., 2020). Indeed, different approaches may yield different metrics for assessing interaction strength, and the resulting mispredictions might affect the reliability of the ACE2 interactions with the spike protein of SARS-CoV-2. Therefore, to find the best metric for assessing ACE2-RBD interactions, we devised and implemented a rigorous computational approach to generate ACE2-RBD protein complex models for 299 ACE2 orthologs, and then benchmarked the predicted dissociation constant (Kd), and change in Gibbs free energy (ΔG) against experimental values (n = 400) retrieved from the PDBbind database. The results revealed that the predicted dissociation constants are highly correlated with experimental values (r = 0.858, P < 0.0001), and that predicted binding affinity of dACE2/hACE2 to RBD is comparable to the experimental binding affinity (Zhang et al., 2021). Together, these results support the robustness and reliability of the adopted approach and findings in this study.

It is of note that the findings are based on predicted propensity (dissociation constants) to bind to SARS-CoV-2 for categorization of vertebrate species into very high, high, medium, and low propensity categories. Intriguingly, the results showed that predicted binding affinity of ACE2 with RBD of SARS-CoV-2 based on the dissociation constants is a better descriptor of species susceptibility to SARS-CoV-2 because all the 46 vertebrate species known to support SARS-CoV-2 infection on the basis of natural and experimental infections were predicted correctly in our study (WHO, 2021). Of note, new world monkey, such as marmosets, was predicted as less susceptible or resistant to SARS-CoV-2 infection in previous studies (Damas et al., 2020; Liu et al., 2021a; Shi et al., 2020), however, in contrast, the present study defined marmoset belonging to medium propensity category (10 times lower predicted binding affinity than that of hACE2). This finding is supported by a recent experimental study in which older marmosets developed mild infection on exposure to SARS-CoV-2 (Singh et al., 2021). Furthermore, high susceptibility of white tailed deer predicted in our study is in line with recent studies demonstrating that white tailed deer are highly susceptible to SARS-CoV-2 infection naturally (Cool et al., 2021; Kuchipudi et al., 2021). Additionally, this study showed a medium propensity for the cattle and pig, which is in line with a previous study showing the efficient entry of SARS-CoV-2 in A549 cells expressing the ACE2 of cattle and pig (Liu et al., 2021a).

Next, we found that the previous studies that had proposed five critical ACE2 interacting residues based on their conservation in known susceptible species (31K/T, 35E/K, 38D/E, 82T/M/N, and 353K) were inconsistent with the comparatively large and diverse ACE2 dataset presented in this study (Shang et al., 2020a; Shang et al., 2020b; Wan et al., 2020). Therefore, based on the sequence alignment of ACE2 interacting residues, we propose six unique residues that could together help in differentiation of susceptible from the resistant species; Susceptible species (27T/I, 30D/E/Q, 82M/T/D/N, 326G/E/R/T, and 352G+353K+354G), resistant species (N326+N330), and 352G+353K+354H/R/Q for reduced susceptibility to SARS-CoV-2 infection. Furthermore, the current approach differs substantially from the previous *in silico* approaches in several aspects: 1) a phylogenetic analysis was conducted for the ACE2 orthologs from 299 vertebrate species to show the existence of considerably high evolutionary diversity across the six vertebrate classes; 2) the ACE2-RBD protein complex-models for 299 vertebrate ACE2 proteins were generated by implementing a robust computational modeling approach for subsequently selecting the best model for downstream processing; 3) the best metric (dissociation constant) was chosen for predicting the binding affinity of ACE2 with the spike protein of SARS-CoV-2 based on the benchmarking with the experimental data; 4) the different ACE2 isoforms were analyzed in a particular bat species to reveal their varied binding affinity to the spike protein of SARS-CoV-2. As a result, we believe that using these approaches allowed us to generate a more realistic representation of species at risk, and species that could potentially serve as the intermediate hosts. Though, the findings suggested many vertebrate species likely to be potential intermediate hosts of SARS-CoV-2, it does not follow that the true intermediate host must be one of them. The list can be narrowed even further by considering the animals’ living conditions, particularly their proximity to human dwellings.

The increasing pieces of evidences have suggested that the binding affinity of ACE2 orthologs from different bat species to the RBD of SARS-CoV-2 differed significantly, implying the existence of diversified susceptibility of bat species to SARS-CoV-2 (Boni et al., 2020; Latinne et al., 2020). The results in the present study are moderately in line with the previous *in silico* studies conducted for the susceptibility prediction of bat species to SARS-CoV-2, because of implementation of different approaches for the prediction of binding affinity. For instance, unlike the previous *in silico* studies, the predictions of bat species susceptibility to SARS-CoV-2 in this study are partially consistent with a recent functional experiment study that utilized 293T cells expressing bat ACE2 orthologues to assess the bat species susceptibility by pseudotyped virus entry assay (Yan et al., 2021). However, this functional experiment also shows disparities with another functional study (Zhou et al., 2021); where the later showed that HeLa cells expressing *Rhinolophus siniscus* ACE2 could serve as an entry receptor for SARS-CoV-2, in contrast the first study. Of note, based on the present study, it is observed that *Rhinolophus siniscus, Rhinolophus affinis*, and *Rhinolophus macrotis* have moderate binding affinity to the spike protein of SARS-CoV-2 as compared to hACE2. These results are consistent with previous functional experiments (Li et al., 2021; Liu et al., 2021b; Zhou et al., 2020). In summary, the findings in the present study show a considerable consistency with the functional experiments than that of previous *in silico* approaches. Moreover, there were also some inconsistencies in binding of RBD to ACE2 of *R. siniscus* in different studies (Liu et al., 2021a; Wu et al., 2020; Yan et al., 2021; Zhou et al., 2020). These inconsistencies are possible owing to the presence of at least 10 ACE2 isoforms in *R. siniscus*, which possess predicted varied binding affinity to the spike protein of SARS-CoV-2. These findings suggested that the ACE2 varied binding affinity owing to different isoforms in a particular bat species could have direct implications in the spillover events on the basis of distribution of these isoforms in different tissues. It would be of interest to delineate the tissue-specific expression of these ACE2 isoforms in future studies.

In conclusion, current findings support the bat origin of SARS-CoV-2 and the involvement of intermediate hosts in virus transmission to humans based on the predicted binding affinity of 299 ACE2 orthologs from various vertebrate species to the spike protein of SARS-CoV-2. It also sheds light on the wide host range and cross-species transmission. Furthermore, SARS-CoV-2 may be far more widespread than previously suggested, emphasizing the importance of intense surveillance programs finding susceptible hosts, particularly their ability to cause zoonosis, in order to prevent future outbreaks.

## Supporting information

The species-wise number of the ACE2 orthologs retrieved from the NCBI protein database are provided in Supplementary Table S1.

Supplementary Table S2 has more information on these protein complexes.

implemented in CPP4 software suite (Supplementary Table S3)

with sequence identities of 96.05-96.86% (Supplementary Table S4). However, the ACE2

The results for all the vertebrate species are provided in Supplementary Table S5

the rest of 8 bat species were predicted to be resistant to SARS-CoV-2 (Supplementary Table S6).

selected vertebrate species are provided in Supplementary Table S7.

## Acknowledgments

The authors thank RIKEN Advanced Center for Computing and Communication (ACCC) for computing resources on the Hokusai BigWaterfall supercomputer. Finally, all of the authors express their appreciation to their respective Institutes.

